# DYRK2 is a ciliary kinase involved in vertebrate Hedgehog signal transduction

**DOI:** 10.1101/2020.08.31.275511

**Authors:** Nicholas Morante, Monika Abedin Sigg, Luke Strauskulage, David R. Raleigh, Jeremy F. Reiter

## Abstract

Primary cilia are organelles specialized for signaling. We previously defined the proteomes of sea urchin and sea anemone cilia to identify ciliary proteins that predate the origin of bilateria. This evolutionary perspective on cilia identified DYRK2, a kinase not been previously implicated in ciliary biology. We found that DYRK2 localizes to cilia and that loss of DYRK2 disrupts ciliary morphology. We also found that DYRK2 participates in ciliary Hh signal transduction, communicating between SMO and GLI transcription factors. Mutation of mouse *Dyrk2* resulted in skeletal defects reminiscent of those caused by loss of *Indian hedgehog* (*Ihh*). Like *Dyrk2* mutations, pharmacological inhibition of DYRK2 dysregulates ciliary length control and attenuates Hedgehog signaling. Thus, DYRK2 is required for ciliary morphology, for Hedgehog signaling *in vitro*, and for skeletal development. We propose that DYRK2 is part of the mechanism that transduces SMO to activate GLI transcription factors within cilia.

## INTRODUCTION

Primary cilia participate in some forms of intercellular communication. One critical signaling pathway transduced via primary cilia is the Hedgehog (HH) pathway. In the absence of HH ligand, its receptor, Patched1 (PTCH1), localizes to the cilium where it represses HH activation by blocking accumulation of SMO at the cilium (Corbit, Aanstad et al. 2005, Rohatgi, Milenkovic et al. 2007). Simulating cells with HH or Smoothened Agonist (SAG), a pharmacological SMO agonist (Chen, Taipale et al. 2002), stimulates SMO accumulation at the cilium. Ciliary SMO activates GLI transcription factors, but the molecular mechanisms by which it does so remains unclear.

The HH signaling pathway has diverse roles in development, including patterning of the neural tube and digits via Sonic hedgehog (SHH) (Hill 1995, Ericson, Briscoe et al. 1997, Sharpe, Lettice et al. 1999) and differentiation and proliferation of chondrocytes via IHH (St-Jacques, Hammerschmidt et al. 1999).

The association of the HH pathway with primary cilia may be more ancient than the origin of chordates as sea urchins require cilia for HH signal transduction (Warner, McCarthy et al. 2014). Sea anemones also possess HH signaling pathway components, although it remains unclear whether they require cilia for HH signal transduction (Matus, Magie et al. 2008). Correlating with the absence of evidence of DYRK2 in their cilia, choanoflagellates do not possess a canonical HH pathway.

To identify ciliary signal transduction components, we previously defined the choanoflagellate, sea anemone and sea urchin ciliary proteomes and identified signaling regulators not previously associated with cilia (Sigg, Menchen et al. 2017). We detected Dual specificity tyrosine-phosphorylation-regulated kinase 2 (DYRK2) in both the sea anemone and sea urchin ciliary proteomes, but not the choanoflagellate ciliary proteome. DYRK2 is a member of a family of kinases conserved from basal eukaryotes to humans and implicated in diverse processes such as E3 ubiquitin ligase complex formation and Nuclear Factor of Activated T-cells (NFAT) sequestration (Maddika and Chen 2009, Aranda, Laguna et al. 2011).

Inspired by its putative localization in cilia and its strong evolutionary conservation, we sought to elucidate the roles of mammalian DYRK2. Here, we show that DYRK2 is required for ciliary morphogenesis, HH signaling, and endochondral ossification, revealing DYRK2 to be an essential regulator of ciliary HH signaling.

## RESULTS AND DISCUSSION

### DYRK2 is an evolutionarily conserved ciliary kinase

Homologs of DYRK2 are present in diverse eukaryotes, and orthologs of mammalian DYRK2 are present in animals, from Trichoplax to humans (Supplementary figure 1A and B). We recently identified DYRK2 as a member of the ciliary proteomes of sea anemones and sea urchins (Figure 1A) (Sigg, Menchen et al. 2017). The detection of DYRK2 protein in the ciliary proteomes of clades evolutionary distant from mammals prompted us to examine its localization in mammalian cells.

**Figure 1:**
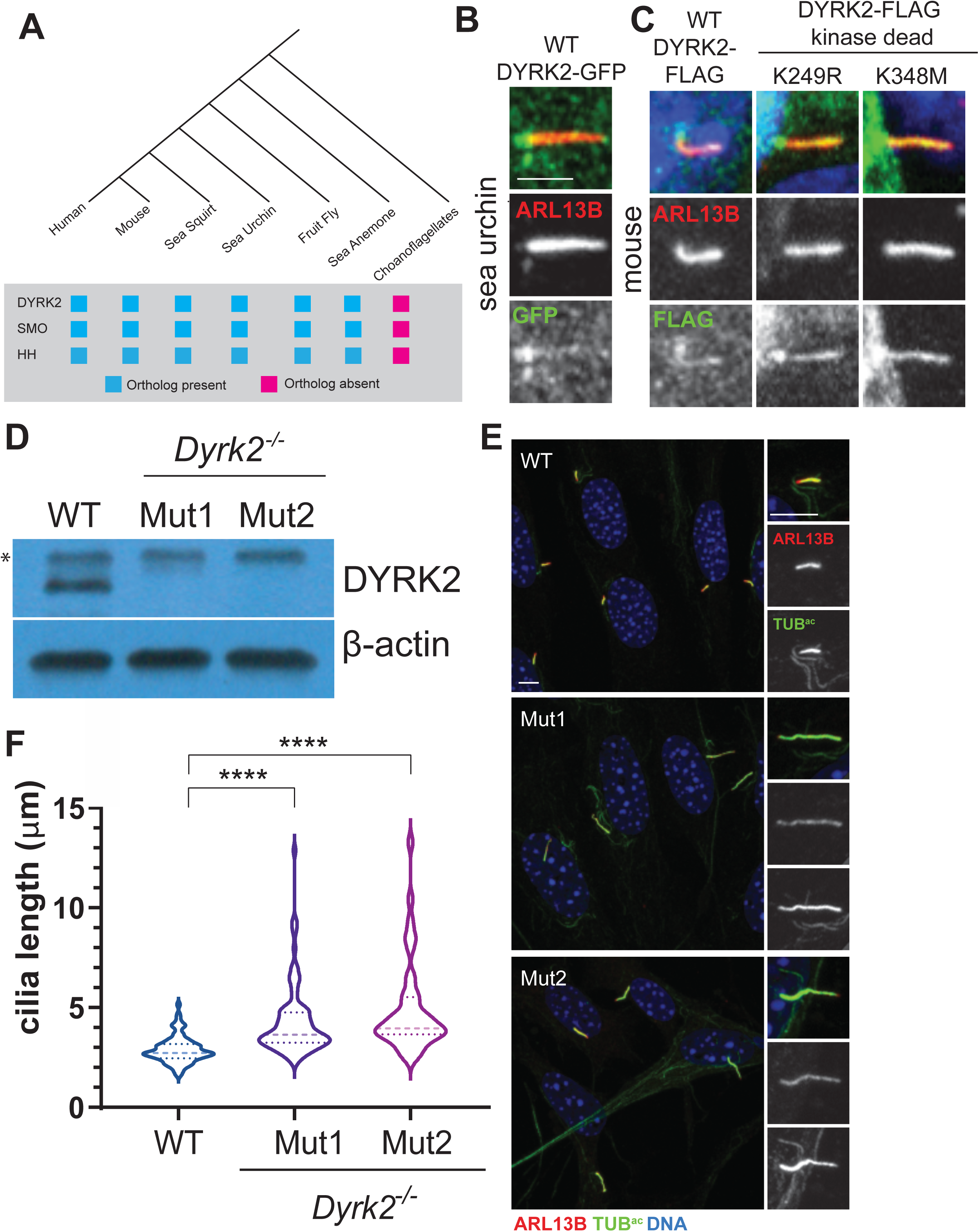
DYRK2 is a conserved ciliary kinase that controls ciliary length. **A)** Phylogeny of DYRK2 and HH and SMO, key components of the HH signaling pathway. **B)** Immunofluoresence imaging of RPE-1 cells expressing DYRK2-GFP. Images depict cells stained for DYRK2-GFP (GFP, green) and cilia (ARL13B, red). **C)** Immunofluorescence imaging of IMCD3 cells expressing mammalian wild type DYR2K-FLAG or versions of DYRK2-FLAG incorporating kinase inactivating mutations K249R or K348M. Images depict cells stained for DYR2K-FLAG (FLAG, green), cilia (ARL13B, red), and nuclei (Hoechst, blue). Scale bar=2.5μm **D)** Immunoblot for DYRK2 protein in wild type NIH/3T3 cells and frameshift mutant lines (Mut1 and Mut2). β-actin immunoblot serves as a loading control. **E)** Immunofluoresence imaging of serum-starved NIH/3T3 cells stained for the ciliary membrane (ARL13B, red), ciliary axoneme (acetylated tubulin [TUB^AC^], green) and nuclei (Hoechst, blue). Scale bar: 5μm **F)** Quantification of ciliary length in wild type, *Dyrk2*-mutant Mut1 and Mut2 NIH/3T3 cells. Dotted lines correspond to the mean and quartiles. A p value less than 0.05 was considered statistically significant and is denoted as follows: ****p<0.0001, ***p<0.001, **p<0.01, *p<0.05, ns= not significant. Significance was assessed using ordinary one-way ANOVA.

To confirm the proteomics indication that DYRK2 can localize to cilia, we expressed a GFP-tagged form of sea urchin DYRK2 in human retinal pigment epithelium (RPE-1) cells and found that it was enriched in primary cilia (Figure 1B). To determine if mammalian DYRK2 also localizes to cilia, we expressed FLAG-tagged mouse DYRK2 in inner medullary collecting duct (IMCD3) cells and found that it similarly localized to primary cilia (Figure 1C). As kinase dead forms of FLAG-tagged DYRK2 also localized to cilia, the serine/threonine kinase activity of mouse DYRK2 is not required for ciliary localization (Figure 1C).

### DYRK2 controls ciliary length

To investigate whether DYRK2 functions in ciliogenesis or ciliary signaling, we used CRISPR-mediated genome editing to mutate *Dyrk2* in mouse NIH/3T3 cells. We created two NIH/3T3 cell lines (referred to as Mut1 and Mut2), each possessing two deletion alleles of *Dyrk2* that alter the reading frame (Supplemental figure 2A). As assessed by immunoblotting for DYRK2, neither Mut1 and Mut2 cell produced detectable levels of DYRK2 (Figure 1D).

**Figure 2:**
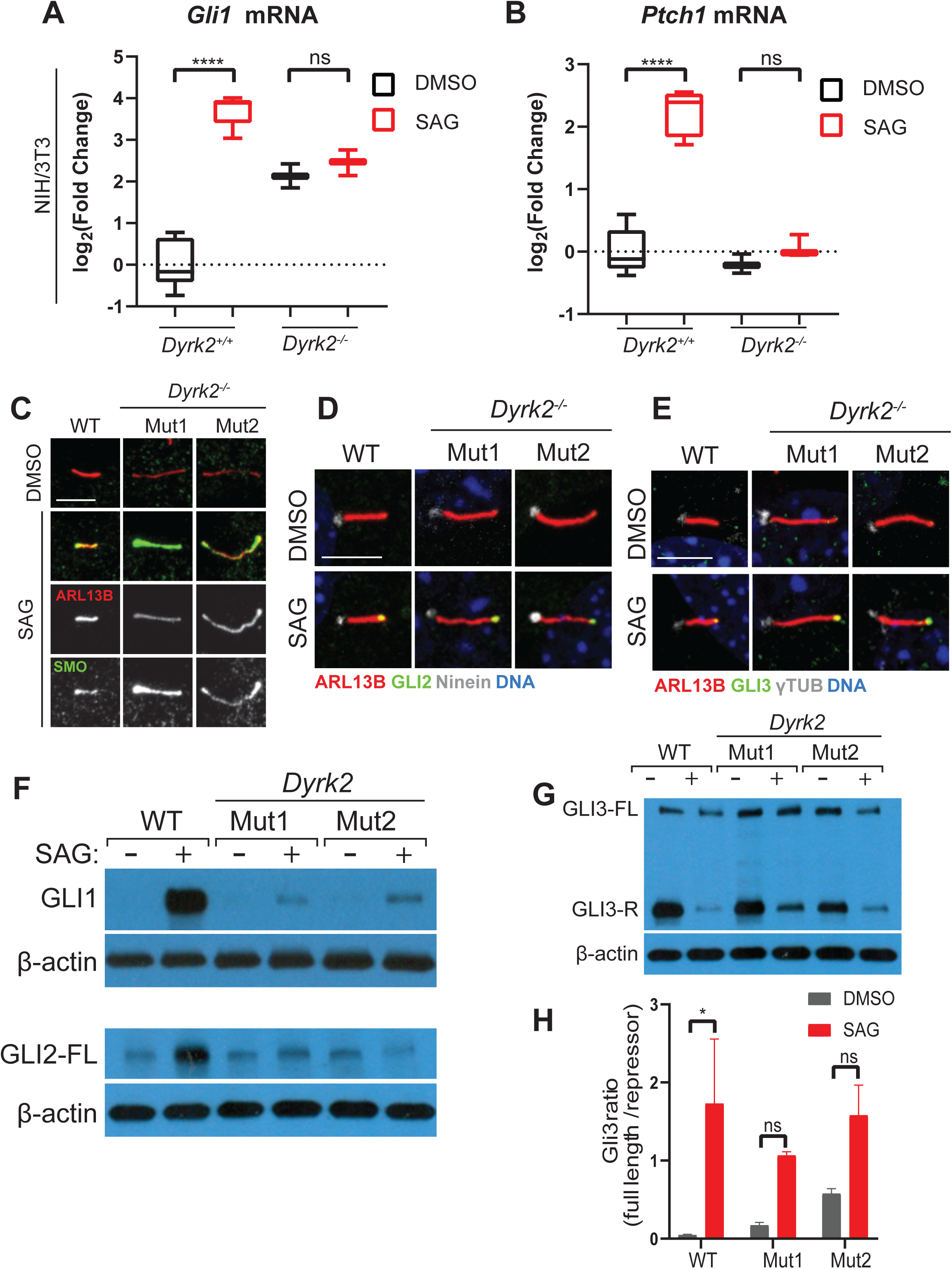
DYRK2 is required for HH signal transduction. **A**,**B)** qRT-PCR quantification of levels of HH targets *Gli1* (A) and *Ptch1* (B) in serum-starved NIH/3T3 cells stimulated with SAG or vehicle control (DMSO). Box and whisker plots indicate the minimum and maximum values, second and third quartiles, and median values. **C-E)** Immunofluorescent images of wild type and mutant NIH/3T3 cell lines. (C) Trafficking of SMO (green) to the cilia, marked with ARL13B (red), under SAG stimulation. (D and E) Trafficking of GLI2 (D) and GLI3 (E) (green) to the ciliary tips in response to SAG stimulation. Basal bodies (gray) are labeled with Ninein (C) or --tubulin (D), cilia labeled with ARL13B (red), nuclei labeled with Hoechst (blue). Scale bar: 5μm. **F**,**G)** Immunoblots for GLI1, GLI2, and GLI3 in wild type and *Dyrk2*-mutant Mut1 and Mut2 cell lines. β-actin immunoblot serves as a loading control. (G) Immunoblot showing full-length GLI3 (FL) and the processed repressor form of GLI3 (R) stimulated with SAG or vehicle control (DMSO). **H)** Quantification of ratio of full-length to repressor forms of GLI3 cells with and without SAG stimulation.

To test if DYRK2 is required for ciliogenesis, Mut1, Mut2 and parental wild-type NIH/3T3 cells were grown to confluency and serum starved to induce ciliogenesis. The cells were stained with antibodies against ciliary membrane component ARL13B (Caspary, Larkins et al. 2007) and ciliary axoneme component acetylated tubulin. *Dyrk2* mutant cells possessed cilia, indicating that DYRK2 is dispensable for ciliogenesis (Figure 1E). Cilia in both *Dyrk2* mutant lines were markedly longer compared to those of the wild-type control cells (Figure 1F). Quantification of cilia length confirmed a mean difference of 1.38 ± 0.237μm (p<0.0001) and 2.04 ± 0.298μm (p<0.0001) between wild-type cilia and mutant lines Mut1 and Mut2, respectively, indicating that DYRK2 restricts ciliary length.

### DYRK2 is required for HH signaling

Components of the HH signal transduction pathway are dispensable for ciliogenesis. As DYRK2 was not required for ciliogenesis, we tested whether it participated in HH signaling. Loss of DYRK2 in NIH/3T3 cells blocked induction of the HH pathway target genes *Gli1* and *Ptch1* by SAG suggesting that, contrary to a previous report (Varjosalo, Björklund et al. 2008), DYRK2 positively regulates the HH pathway (Figure 2A and B).

Because DYRK2 is required for activation of the HH pathway by SAG, a direct SMO agonist, DYRK2 acts either at the level of SMO or downstream of it. To determine if DYRK2 regulates accumulation of SMO at the cilium, we analyzed the localization of endogenous SMO in wild type *Dyrk2* mutant NIH/3T3 cells. As in wild type NIH/3T3 cells, SMO was absent from cilia in unstimulated *Dyrk2* mutant cells and became enriched in cilia upon stimulation with SAG, suggesting that DYRK2 is required for HH signal transduction downstream of the accumulation of SMO within cilia (Figure 2C).

Ciliary SMO enriches the localization of the downstream transcriptional effectors GLI2 and GLI3 at the ciliary tip (Haycraft, Banizs et al. 2005, Kim, Kato et al. 2009, Wen, Lai et al.2010). As in wild type NIH/3T3 cells, GLI2 and GLI3 localized to the tips of cilia and became enriched upon stimulation with SAG in *Dyrk2* mutant cells (Figure 2D and E)

GLI2 and GLI3 can act as either transcriptional activators or repressors, whereas GLI1 is a dedicated activator (Dai, Akimaru et al. 1999, Sasaki, Nishizaki et al. 1999). To determine if DYRK2 regulates GLI protein levels, we immunoblotted lysates from wild type and *Dyrk2* mutant cells that were either unstimulated or stimulated with SAG for GLI1, GLI2 and GLI3. In wild type cells, activation of the HH pathway increased levels of GLI2, whereas in the absence of DYRK2, pathway stimulation had no effect on GLI2 levels (Figure 2F). Similarly, in the absence of DYRK2, pathway activation had little effect on GLI1 levels (Figure 2F).

In the absence of HH stimulation, GLI3 is proteolytically processed into a truncated repressor form, and the presence of HH suppresses this processing (Wang, Fallon et al. 2000). To determine if DYRK2 regulates GLI3 processing, we examined how pathway stimulation affected the levels of full length GLI3 and truncated GLI3 in control and *Dyrk2* mutant cells. DYRK2 was essential for the full reduction in GLI3 processing upon pathway stimulation with SAG (Figure 2G and H). As the accumulation of GLI2 and GLI3 at ciliary tips is independent of DYRK2, and the regulation of GLI1 and GLI2 levels and GLI3 processing is dependent on DYRK2, we propose that DYRK2 is required for the HH-regulated activation of GLI proteins at the ciliary tip.

### DYRK2 is required for skeletal development

To explore the in vivo function of DYRK2, we generated a mutation in mouse *Dyrk2* by CRISPR/Cas9-mediated deletion of the first two exons of *Dyrk2*, including the TATA box and initiator codon. (Figure 3A). In agreement with the results obtained using mutant NIH/3T3 cells, MEFs obtained from *Dyrk2*^-/-^ mice failed to induce *Gli1* and *Ptch1* in response to stimulation with SAG (Figure 3B and C).

**Figure 3:**
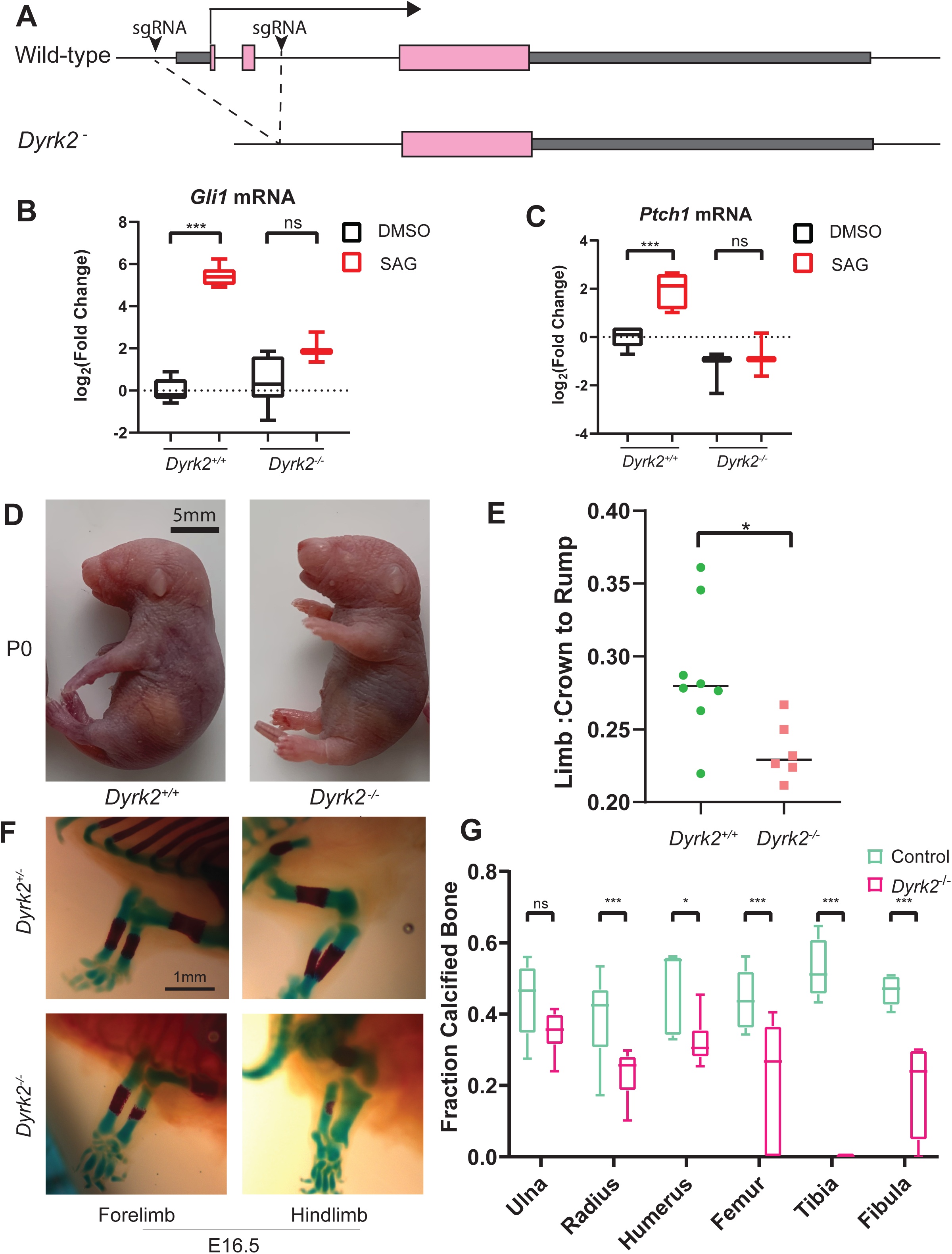
DYRK2 is essential for mouse limb development. **A)** Diagram of the CRISPR/Cas9-mediated excision of the first two exons of *Dyrk2* in mouse. The bent arrow indicates the direction of transcription from the transcriptional start site in exon 1 while the small arrowheads represent Cas9/sgRNA target sites. Pink boxes represent coding regions of the *Dyrk2* gene and grey boxes indicate the 5’ and 3’ UTRs. The dashed lines indicate the region deleted via CRISPR/Cas9 mutagenesis. **B**,**C)** Box and whisker plots of qRT-PCR of *Dyrk2*^-/-^ and littermate control MEF lines treated with SAG or vehicle control (DMSO) for HH target genes *Gli1* (B) and *Ptch1* (C). **D)** Gross images of neonatal (P0) *Dyrk2*^-/-^ and littermate control mice. Scale bar: 5mm. **E)** Quantification of the ratio of the length of the limb (stylopod plus zeugopod) to the crown-rump length of *Dyrk2*^-/-^ and littermate control mice. Significance was assessed using two-tailed t-test. **F)** Wholemount images of alizarin red and alcian blue-stained bone preparations of limbs from E16.5 *Dyrk2*^-/-^ and *Dyrk2*^-/+^ littermate control mice. **G)** Quantitation of endochondral ossification, measured by the ratio of the extent of alizarin red staining to the length of the bone. Significance was calculated using multiple unpaired t-tests with the Holm-Šídák test.

No *Dyrk2*^-/-^ pups survived past two days of birth, indicating that DYRK2 is essential for perinatal viability. Truncation of the limbs was evident in *Dyrk2*^-/-^ pups at P0 (Figure 3D). The ratio of the length of *Dyrk2*^-/-^ limbs compared to the crown-to-rump measurement was significantly reduced as compared to control littermates (Figure 3E). Furthermore, rare instances of supernumerary digits were observed in *Dyrk2*^-/-^ limbs (Supplemental figure 2A, 5/37 mutants assessed).

The reduction in limb elongation observed in *Dyrk2* mutants is reminiscent of that caused by *Ihh* loss of function mutations (St-Jacques, Hammerschmidt et al. 1999). In *Ihh* mutant mice, defective chondrocyte differentiation causes defects in calcification of the long bones. We therefore assessed bone ossification of embryonic and neonatal *Dyrk2* mutants utilizing alizarin red and alcian blue staining to measure calcified bone and cartilage, respectively (Figure 3F). There was a decrease in ossification in *Dyrk2*^-/-^ long bones at embryonic day (E) 16.5 relative to wild-type littermate controls (Figure 3G). Neonatal *Dyrk2* mutants also exhibited reduced calcification of long bones (Supplemental figure 2B,C). Thus, DYRK2, like IHH, is required for long bone growth and calcification.

### DYRK2 is a pharmacologically tractable candidate for the treatment of HH pathway-associated cancers

Misactivation of HH signaling is critical for the formation of some cancers, including a subset of medulloblastomas (Goodrich, Milenković et al. 1997, Vorechovský, Tingby et al. 1997, Vôrechovský, Undén et al. 1997, Zurawel, Allen et al. 2000, Taylor, Liu et al. 2002). To test whether DYRK2 might function in oncogenic HH signaling, we examined *Dyrk2* expression in a mouse model of HH-associated medulloblastoma caused by a constitutively active form of SMO (Xie, Murone et al. 1998, Mao, Ligon et al. 2006, Schüller, Heine et al. 2008). *Dyrk2* expression and DYRK2 protein levels were increased in medulloblastomas (Figure 4A and B).

**Figure 4:**
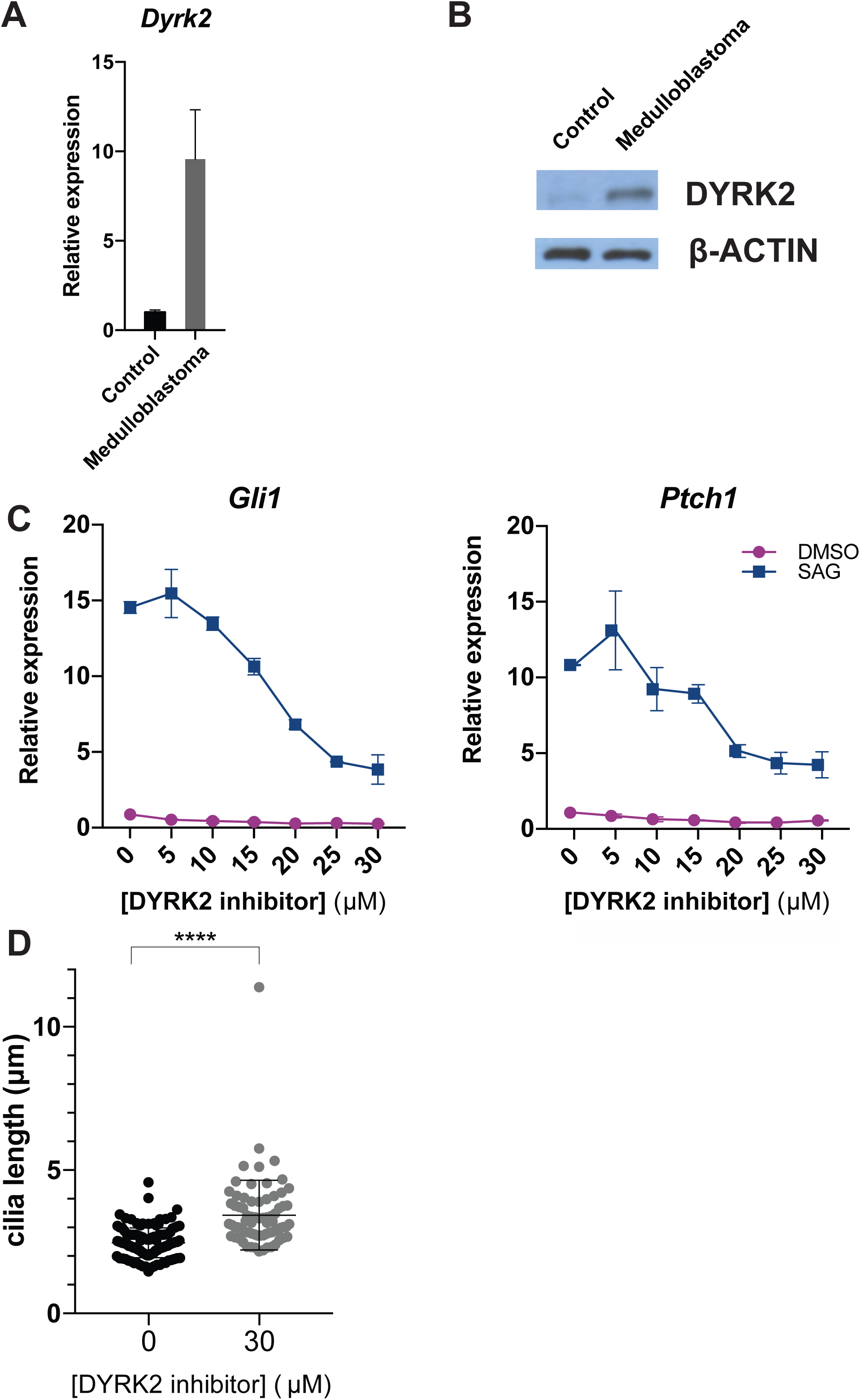
DYRK2 is a potentially druggable target in HH pathway-associated cancers. **A)** PCR measurement of *Dyrk2* expression in control (*SmoM2*^cond/wt^) mouse cerebellum tissue and medulloblastoma tumor (*Math1-Cre SmoM2*^cond/wt^) tissue. Error bars represent SEM, 4 mice per condition. **B)** Immunoblot for DYRK2 of control cerebellar and medulloblastoma tumor lysates. β-actin immunoblot serves as a loading control. **C**,**D)** qRT-PCR measurement of *Gli1* and *Ptch1* expression in MEFs in the presence of the indicated concentrations of compound 24 (Schmitt et al., 2014), a DYRK2 inhibitor, without stimulation (DMSO) or after stimulation with SAG. The DYRK2 inhibitor blocks HH pathway activity in a concentration dependent manner. The experiment was repeated 3 times and the data shown are representative from 1 experiment. Error bars show SD of 2 technical replicates. **E)** Quantification of ciliary length in NIH/3T3 cells treated with DYRK2 inhibitor.

Pharmacological inhibition of the HH pathway is an effective therapy for some forms of HH pathway-associated cancers (Von Hoff, LoRusso et al. 2009). Therefore, we tested whether a pharmacological inhibitor of DYRK2 (Schmitt, Kail et al. 2014) could inhibit HH signaling in mouse embryonic fibroblasts (MEFs). DYRK2 inhibitor had no effect on SMO localization to cilia, but blocked SAG-dependent *Gli1* and *Ptch1* induction in a concentration dependent manner (Figure 4C). Pharmacological inhibition of other DYRK family members did not block HH signaling (Supplemental figure 3A). Moreover, DYRK2 inhibitor caused the MEFs to form longer cilia (Figure 4D). Thus, a small molecule inhibitor of DYRK2 reproduced the signaling and ciliary morphology effects of loss of function mutations in *Dyrk2*. These results reveal that evolutionary proteomics can help identify novel functional components of an organelle, in this case a ciliary-localized kinase critical for the activation of the HH signaling pathway. We anticipate that future analysis of ciliome members will uncover still other HH pathway regulators.

We predicted that proteins shared between the sea urchin and sea anemone ciliary proteomes, such as DYRK2, would function in ciliary biology. We further proposed that ciliary proteins conserved specifically among animals that employ HH signaling may function in HH signaling. In this work, we found that one such protein, DYRK2, localizes to cilia and determined, using a combination of CRISPR/Cas9-mediated knockout and pharmacological inhibition in both mammalian cells and mice, that DYRK2 positively regulates the HH pathway downstream of SMO.

Little is known about how information is communicated between vertebrate SMO and the GLI transcriptional effectors of the HH pathway. The identification of DYRK2 as epistatic to SMO ciliary accumulation suggests that this kinase could function at this important and mysterious step in HH signal transduction. Moreover, the requirement for DYRK2, not in ciliary tip accumulation of GLI2 and GLI3, but in increasing GLI1 and GLI2 protein levels and in GLI3 processing, further points to DYRK2 as an important mediator of information between SMO and GLI proteins.

In addition to its requirement for HH signaling, DYRK2 limits the length of ciliary microtubules. Another member of the DYRK protein family, DYRK1A, phosphorylates β-Tubulin to regulate cytoplasmic microtubule dynamics (Ori-McKenney, McKenney et al. 2016). It will be interesting to test whether DYRK2 has an analogous role in the regulation of ciliary microtubules.

As DYRK2 is required for HH signal transduction in NIH/3T3 cells, it was surprising that *Dyrk2* mutant embryos exhibit phenotypes that are reflective of moderate attenuation of HH signaling. For example, polydactyly is pronounced in embryos with strong defects in mouse embryos with strongly defective ciliary signaling, such as *Tctn1* mutants (Garcia-Gonzalo, Corbit et al. 2011). Similarly, mutation of genes encoding some key components of the HH signal transduction pathway, such as *Ptch1* or *Kif7*, cause completely penetrant polydactyly (Liem, He et al. 2009). In contrast, *Dyrk2* mutants, displayed weakly penetrant polydactyly and bone growth and ossification defects. The *Dyrk2* mutant phenotypes suggest tissue specific requirements or a modulatory role rather than a global requirement for HH signaling. It is possible that other dual-specificity kinases partially compensate for loss of DYRK2 in some cell types during development.

The defects in long bone growth and endochondral ossification of *Dyrk2* mutants is reminiscent of those defects caused by loss of *Ihh*. Like *Dyrk2* mutants, *Ihh* mutants display attenuated long bone growth and endochondral ossification (St-Jacques, Hammerschmidt et al. 1999). IHH is essential for the differentiation and proliferation of chondrocytes. The similarity of these phenotypes leads us to propose that DYRK2 transduces IHH signals in chondrocytes.

DYRK2 is overexpressed in a mouse model for medulloblastoma. The efficacy of *in vitro* inhibition of DYRK2 and its role in enabling HH signaling in certain contexts raises the possibility that pharmacologically inhibiting DYRK2 may be beneficial in the treatment of HH pathway-associated cancers. As resistance to existing HH pathway inhibitors, such as vismodegib, can occur at the level of SMO (Yauch, Dijkgraaf et al. 2009, Brinkhuizen, Reinders et al. 2014, Pricl, Cortelazzi et al. 2015, Sharpe, Pau et al. 2015), inhibiting the pathway downstream of SMO at the level of DYRK2 may be beneficial in vismodegib-resistant cancers. Additionally, inhibition of DYRK2 may synergize with SMO inhibition. Thus, pharmacological inhibition of DYRK2 may be a viable strategy for treating HH pathway-associated cancers such as medulloblastoma and basal cell carcinoma.

## METHODS

### CRISPR/Cas9-mediated mutagenesis

CRISPR-mediated *Dyrk2* mutations were generated as described previously (Ran, Hsu et al. 2013). Briefly, NIH/3T3-FlpIn cells were transfected with pSpCas9(BB)-2A-Puro (PX458) containing one of two small guide sequences to target *Dyrk2* (see Table S4 for primer sequences). 24 hours after transfection, selection media containing 300ng/ml puromycin (InvivoGen) was applied to the cells for 48 hours. The cells were then cultured at low density and individual clones were isolated. To identify clones with mutations in *Dyrk2*, genomic DNA was extracted and primers specific to sequence flanking the small guide target sites were used to PCR amplify a 755bp portion of the *Dyrk2* gene. The amplicons were sequenced by Sanger sequencing to identify mutations.

### Generation of Mouse Embryonic Fibroblast lines

Isolation of MEF lines are adapted from (Jozefczuk, Drews et al. 2012). Briefly, pregnant mice at E15.5 were sacrificed, uterine horns were dissected and dipped in 70%ethanol and placed in ice-cold Delbecco’s phosphate buffered saline. Embryos were extracted from the uterine horns, organs and heads were removed and the embryos were finely minced followed by digestion with Kunitz DNAse and .05%Trypsin-EDTA. The resultant cell suspension was plated and grown to confluency in DMEM containing penicillin and streptomycin solution (Gibco). Cells were immortalized with SV40 virus and maintained.

### Alignments and dendrograms

Alignment generated with ClustalW and Boxplot (Madeira, Park et al. 2019). Phylogenetic tree of the above sequences, calculated with NGPhylogeny.fr (Dereeper, Guignon et al. 2008, Lemoine, Correia et al. 2019). The NCBI accession numbers of the sequences were used in the alignments are as follows: NP_006473.2, NP_001014412.1, NP_001038298.1, XP_030856015.1, NP_001033810.1, XP_001638207.2,XP_002108931.1:14-400, XP_004990443.1, XP_015137468.1, XP_003224671.2, XP_030115200.1, and XP_026694893.1:34-604.

### Mice

The *Dyrk2*^em1(IMPC)Ics^ (*Dyrk2*^-^) mouse allele was generated on the C57BL/6N background by the International Mouse Phenotyping Consortium by CRISPR/Cas9-mediated deletion of the proximal promoter and exons 1 and 2. *B6*.*Cg-Tg(Atoh1-cre)*^*1Bfri*/J^ (*Math1-Cre*), *GT(ROSA)26*^*Sortm1(Smo/EYFP)Amc*/J^ (*SmoM2*^*cond*^) and *Ptch1*^*tm1Mps*/J^ (*Ptch1*^*c*^) alleles were obtained from The Jackson Laboratory (Bar Harbor, ME). *Math1-Cre SmoM2*^*cond*^ animals were monitored for survival until death or protocol-defined neurologic endpoints including hydrocephalus or ataxia.

### Antibodies

Primary antibodies used were anti-β-actin (60008-I-Ig, ProteinTech, Rosemont, IL and 907 ab8227, Abcam, Cambridge, MA), anti-acetylated α-tubulin (clone 6-11B-1, Sigma-Aldrich, St. Louis, IL), anti-β-tubulin (E7, Developmental Studies Hybridoma Bank, Iowa City, Iowa), anti-GFP (ab290, Abcam and sc-9996, Santa Cruz Biotechnology, Inc., Dallas, Texas), anti-FLAG (clone M2, Sigma-Aldrich), anti-DYRK2 (8143, Cell Signaling, Danvers, MA), anti-GLI1 (L42B10, Cell Signaling), anti-GLI2 for immunofluorescence (gift from J. Eggenshwiler (Cho, Ko et al. 2008)), anti-GLI2 for immunoblot (clone 1H6, gift from S. Scales (Wen, Lai et al. 2010)), anti-GLI3 (clone 6F5, gift from S. Scales (Wen, Lai et al. 2010)), anti-SMO (ab38686, Abcam), and anti-ARL13B (clone N295B/66, Neuromab, Davis, CA and 17711-1-AP, ProteinTech), anti-Ninein (gift from J. Sillibourne, (Delgehyr, Sillibourne et al. 2005)), and anti---tubulin (V-20, Santa Cruz Biotechnology).

### Ciliary length quantification

To quantify cilia length in the *Dyrk2* mutant and control cell lines, a freehand line was drawn along the length of each cilium using FIJI software corresponding to the extent of ARL13B immunofluorescent signal and the length of the line was measured. For WT and Mut1, data from 3 experiments was compiled, 3 images per cell line. For Mut2, data from 2 experiments was used (WT n=82, Mut1 n=91, Mut2 n=41). For control and DYRK2 inhibitor treated MEFs, measurements were collected as described above from 1 experiment (untreated n=125, treated n=76). The data was plotted using Prism 7 (Graph Pad).

### Immunoblotting

Mammalian cells were lysed using RIPA buffer (50mM Tris, pH 7.4, 150mM NaCl, 1%NP-40, 0.5%sodium deoxycholate) with protease inhibitors (Calbiochem, Billerica, MA). Protein samples were separated on 4–15%gradient TGX precast gels (Bio-Rad, Hercules, CA). Protein was transferred to a nitrocellulose membrane (Whattman, Pittsburgh, PA). Membranes were blocked and antibodies were diluted in 5%non-fat milk in TBST (50mM Tris, pH 7.6, 150mM NaCl, 0.1%Tween 20) and analyzed using ECL Lightening Plus (Perkin–Elmer, Waltham, MA).

### Quantitative RT-PCR

RNA was extract from cells using the RNeasy Mini or Micro Kit (Qiagen, Hilden, Germany) and cDNA was synthesized using iScript cDNA Synthesis Kit (BioRad). Quantitative PCR was performed using EXPRESS SYBR GreenER with Premixed ROX (Invitrogen) and the 7900HT Fast Real-Time PCR System (ThermoFisher Scientific). Relative expression was calculated using the delta-delta CT method (Livak and Schmittgen 2001). For *Dyrk2, Glyceraldehyde 3-phosphate dehydrogenase* (*Gapdh*) was used as the control for normalization. See Table S1 for primer sequences. The data were analyzed using Microsoft Excel and plotted using Prism7.

### Stimulation of cilia formation, Hh pathway activation and DYRK inhibition

To induce cilia formation in confluent NIH/3T3 cells, cell culture medium was replaced with OptiMEM (CCF, UCSF) for at least 24 hours. To activate the Hh pathway, OptiMEM was supplemented with 1μM SAG (EMD-Millipore, Billerica, MA) for 24 hours.

To inhibit DYRK2, MEFs plated to confluency were treated in cell culture medium with the DYRK2 inhibitor compound 24 (Schmitt, Kail et al. 2014) (gift from M. Engel) for 24 hours. The culture medium was then replaced with OptiMEM containing compound 24 with or without 40μM SAG and incubated for 24 hours. DYRK1A was inhibited using 5μM harmine (R&D Systems) and DYRK1B was inhibited using 5μM AZ191 (Tocris Bioscience, Bristol, United Kingdom) as described for compound 24.

### Bone preparations

Embryonic and neonatal mice were euthanized, fixed overnight in 4%PFA/PBS, and excess tissue was removed before dehydrating in ethanol. The alizarin red and alcian blue staining was performed as described in (Rigueur and Lyons 2014) followed by clearing in glycerol.

### Expression and *in situ* plasmid creation

To construct the GFP-tagged expression plasmids for *S. purpuratus* DYRK2, gene specific cDNA was made from RNA of gastrula stage embryos using AccuScript (Agilent) or SuperScriptIII (Thermo Fisher Scientific), PCR amplified using primers containing the SpeI and AsiSI cut sites and TOPO-TA cloned into PCR2.1 (Invitrogen) and inserted into the pCS2+8CeGFP (Gökirmak et al., 2012).

*M. musculus Dyrk2* cDNA was cloned into pcDNA4/TO-C-3xFLAG using the In-fusion HD cloning system (Clontech). *Dyrk2* cDNA clone #6808145 was purchased (ThermoFisher Scientific). Site directed mutagenesis was performed using the QuikChange Lightning Kit (Agilent) to generate the DYRK2 kinase dead constructs. All primer sequences are listed in Table S1.

## FIGURE LEGENDS

**Figure S1: Evolutionary conservation of Dyrk2**

A) Alignment of protein sequences of human (*H. sapiens*), mouse (*M. musculus*), zebrafish (*D. rerio*), sea urchin (*S. pupuratus*), sea squirt (*C. intestinalis*), sea anemone (*N. vectesis*), placazoan (*T. adhaerens*), fruit fly (*D. melanogaster*), chicken (*G. gallus*), anole (*A. carolinensis*), and zebrafinch (*T. guttata*), orthologs and closest homologous type II DYRK kinases. B) Dendrogram of the above sequences. Note that only partial *T. adhaerens* protein data is available and was excluded from dendrogram. C) Schematic of CRISPR mutations generated in NIH/3T3 cells. Mutations create frameshifts in the third exon.

**Figure S2: DYRK2 inhibits digit formation and endochondral ossification**

**A)** Wholemount images of forelimbs from a wild-type and mutant exhibiting preaxial polydactyly, stained with alizarin red and alcian blue. Scale bar: 1mm **B)** Wholemount images of alizarin red and alcian blue-stained bone preparations of limbs from P0 *Dyrk2*^-/-^ and *Dyrk2*^-/+^ littermate control mice. **C)** Quantitation of endochondral ossification, measured by the ratio of the extent of alizarin red staining to the length of the bone. Significance was calculated using multiple unpaired t-tests with the Holm-Šídák test.

**Figure S3: Pharmacological inhibition of DYRK2 recapitulates mutant phenotype**

**A)** qRT-PCR for *Ptch1* and *Gli1* of NIH/3T3 cells treated with pharmacological inhibitors of DYRK1A or DYRK1B.

**Table S1:**
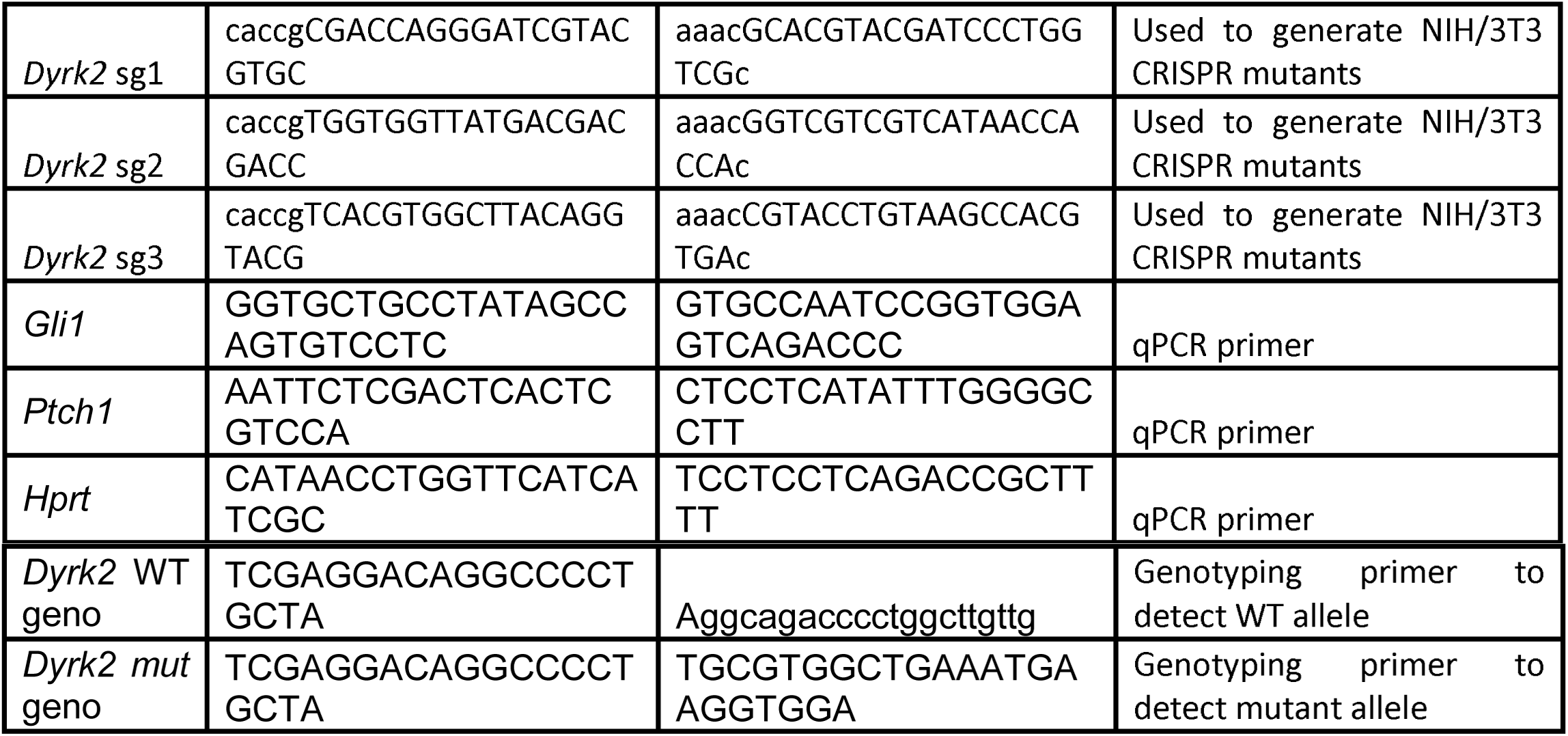
Oligonucleotides Used in Study.

